# Visualizing and sonifying neurodata (ViSoND) for enhanced observation

**DOI:** 10.64898/2026.03.21.713430

**Authors:** Leah Blankenship, Scott C. Sterrett, Dylan M. Martins, Teresa M. Findley, Elliot Abe, Philip R.L. Parker, Cristopher M. Niell, Matthew C. Smear

## Abstract

Neuroscience needs observation. Observation lets us evaluate data quality, judge whether models are biologically realistic, and generate new hypotheses. However, high-dimensional behavioral and neural data are too complex to be easily displayed and eye-tested. Computational methods can reduce the dimensionality of data and reveal statistically robust dynamical structure but often yield results that are difficult to relate back to the underlying biology. In addition, the choice of what parameters to quantify may not capture unexpectedly relevant aspects of the data. To supplement quantification with enhanced qualitative observation, we developed Visualization and Sonification of NeuroData (ViSoND), an open-source approach for displaying multiple data streams using video and sonification. Sonification is nothing new to neuroscience. Scientists have sonified their physiological preparations since Lord Adrian’s earliest recordings. We extend this tradition by mapping multiple physiological datastreams to musical notes using MIDI. Synchronizing MIDI to video provides an opportunity to watch an animal’s movement while listening to physiological signals such as action potentials. Here we provide two demonstrations of this approach. First, we used ViSoND to interpret behavioral structure revealed by a computational model trained on the breathing rhythms of freely behaving mice. Second, ViSoND revealed patterns of neural activity in mouse visual cortex corresponding to eye blinks, events that were previously filtered out of analysis. These use cases show that ViSoND can supplement quantitative rigor with observational interpretability. Additionally, ViSoND provides an accessible way to display data which may broaden the audience for communication of neuroscientific findings.

## Introduction

Technical advances have inspired a resurgence of investigation of complex behavior. Historically, behavior has been studied through observations in laboratory and natural environments, with hypotheses built on human-perceptible patterns (Benzer, 1973; Bullock, 1973; Crowcroft, 1966; Gallistel, 1982; Lorenz, 1978; Muybridge, 1891; Konishi, 1973; Sperry, 1957; Tolman, 1932).

However, human perception and intuition can be overwhelmed by the rich dynamics and high dimensionality of datasets produced by modern neuroethological tools. Therefore, the field has shifted away from observation towards computational models to describe behavior (Bialek, 2022; Branson et al., 2009; Datta et al., 2019; Dennis et al., 2021; Egnor & Branson, 2015; Kennedy, 2022; Mathis & Mathis, 2020; Miller et al., 2022). However, these models can be difficult to interpret biologically, hindering their ability to generate insights that inspire new hypotheses and advance understanding. Importantly, dimensionality reduction and state space modeling necessitate reductionist choices which can lead to loss of key features in the data that the researcher doesn’t already expect (Krakauer et al., 2017; Levins & Lewontin, 1987). To help supplement these computational tools, we have developed an approach to enhance observational investigation of simultaneously acquired behavioral and physiological data. Here, we show how to sonify recordings using a synchronized video and sound format.

Sonification of neurodata has been in use since the inception of neurophysiology by Lord Adrian, who published a circuit diagram for sending recordings to a loudspeaker in one of his first papers (Fig 1A). He also pressed a phonograph record for a demonstration during his Nobel lecture (Adrian,1932). Adrian identified two major benefits of listening to neural recordings. First, human vision is foveal – we can only look in one place at one time. Listening to the physiological signals allows the experimenter to keep their eyes on the preparation. Secondly, he realized that the pattern recognition capabilities of vision and audition are different: “The individual rhythms can often be followed much more easily, … for the ear can pick out … slight differences in intensity and quality which are hard to detect in the complex electrometer record” (Adrian & Bronk, 1928). Electrophysiologists have been listening to brain activity as clicks, pops, and buzzes ever since.

**Figure 1.**
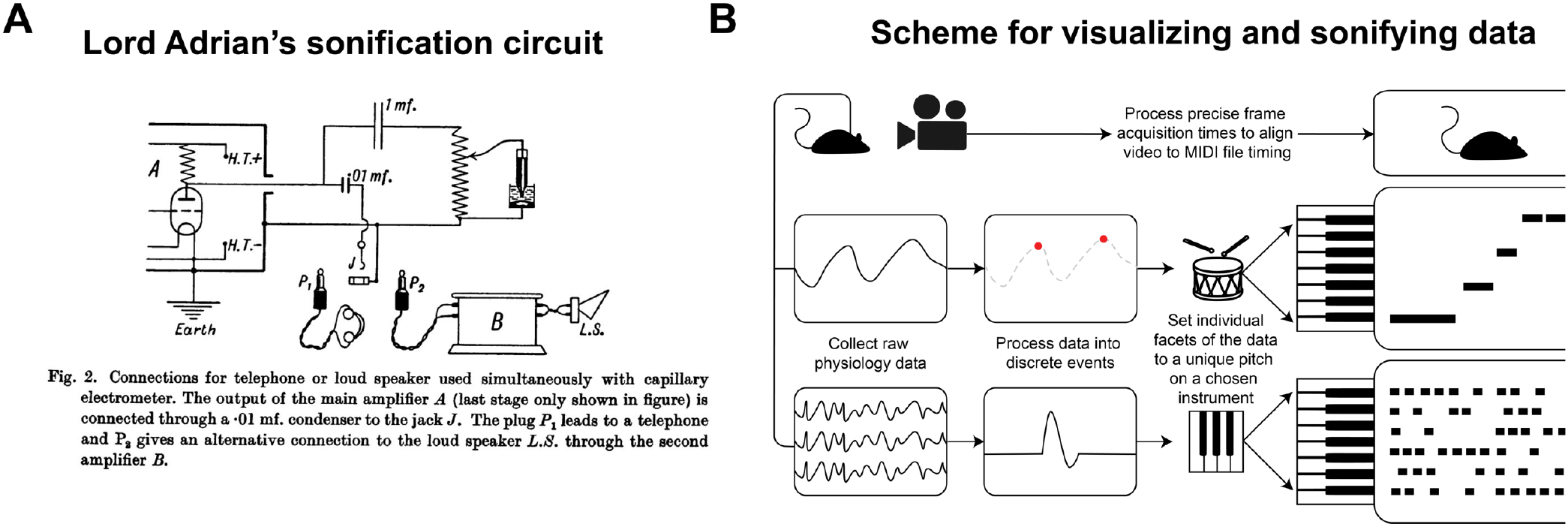
Sonification of neural data. (A) Lord Adrian’s circuit diagram for sonification of his experimental preparation for recording from the phrenic nerve (Adrian and Bronk, 1928). (B) Diagram of the pipeline for capturing, visualizing, and sonifiying multiple streams of data for simultaneous observation.

**Figure 2.**
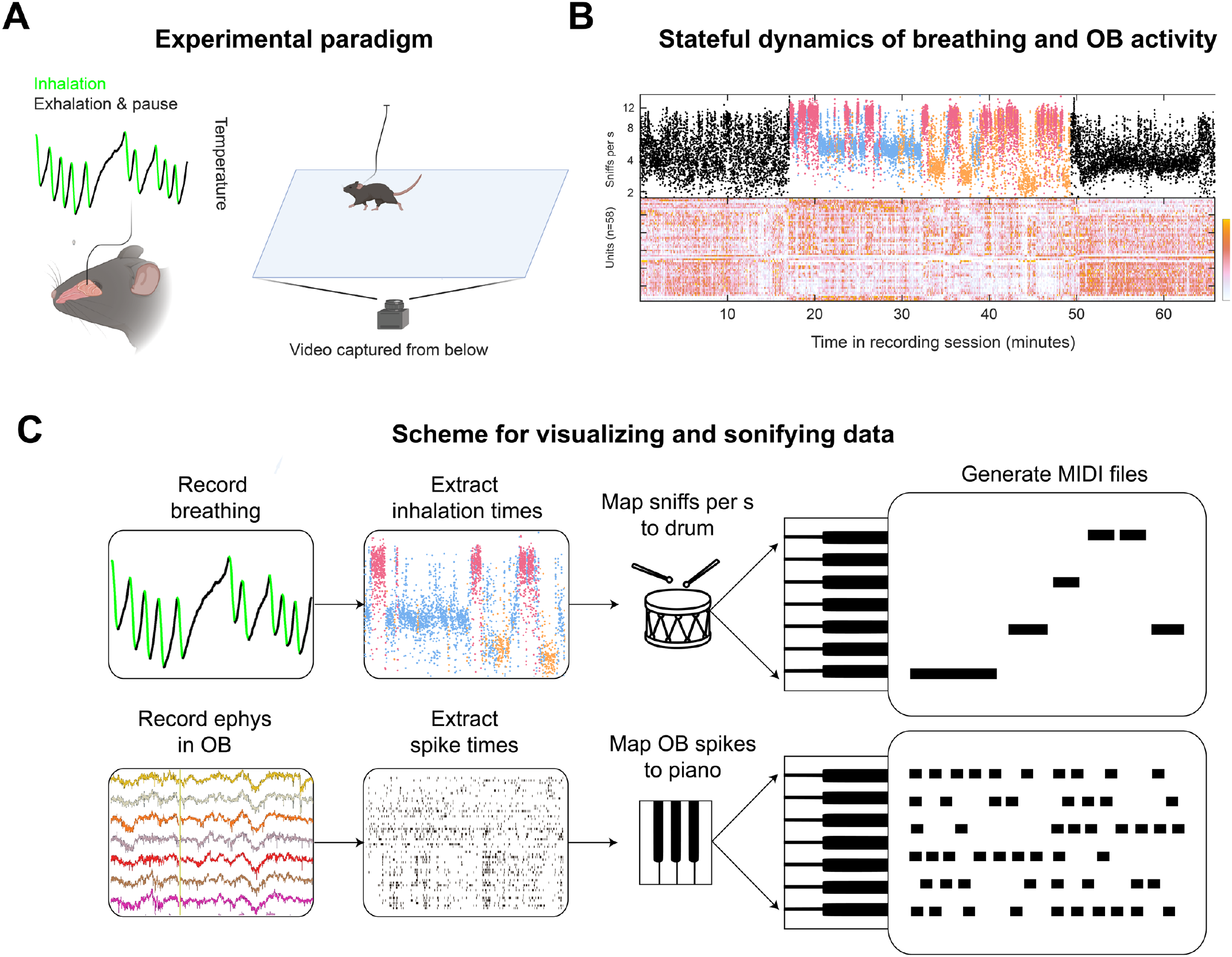
A breathing rhythm associated with grooming (accompanies Video 1). (A) Diagram of experiment with simultaneous recording of spontaneous behavior, olfactory bulb (OB) activity, and respiration with minimally invasive thermal probe. (B) Example data from one session. Colors represent rhythmic states classified by a Hidden Markov Model fit to breathing and pose tracking. (C) Diagram of ViSoND pipeline for the experiment. Data in A and B adapted from Sterrett et al., 2025.

While classical neurophysiologists usually recorded from one neuron at a time, modern multielectrode recordings enable simultaneous recordings from up to thousands of neurons. Even with a small number of neurons, keeping track of which neuron is doing what quickly becomes impossible in a direct sonification of the raw recording. Which pop goes with which cell? To overcome this obstacle, we use pitch to map multiple neural channels, analogous to how anatomists use pseudocolor. It is standard practice to uniquely identify the anatomical processes of different cells with different colors. We use sound to uniquely identify the physiological signals of different cells with different pitches.

To implement this sonification, we render physiological data to Musical Instrument Digital Interface (MIDI) format, which has been the industry standard for controlling electronic music devices for decades. MIDI functions as data instructions that can control hardware or software instruments to play sequences of notes. Therefore, it is a suitable format with which to represent and sonify sequences of any kind of discrete event, of which neuronal spikes provide one example. Importantly, MIDI is well-established and pervasive in the work of professional and hobbyist musicians, so there is a thriving market which has motivated software companies to develop robust, easy-to-use digital audio workstations. This circumvents the need for scientists to develop their own tools for visualizing and sonifying data. It is easy to plug and play ViSoND files into software written by professional developers.

Here, we provide basic instructions to combine our open source ViSoND with two different methods of simultaneous visualization. MIDI files can either be simply played alongside recorded video for free in VLC or for deeper examination, files can be browsed in Ableton Live.

## Methods

### Neural data collection and initial processing

In the Smear lab, we measured breathing with a thermistor placed outside the nasal respiratory epithelium (Findley et al., 2021; McAfee et al., 2016), recorded video with a camera placed beneath the experimental arena, and recorded neural activity using a chronically implanted 64 multi-shank silicon probe in olfactory bulb (Rafilson et al., 2025; Sterrett et al., 2025).

In the Niell lab, pupil orientation was measured from a head-mounted miniature camera, head and body movement was measured from a head-mounted inertial measurement unit (IMU), and single-unit electrophysiology was performed using a chronically implanted 64- or 128-channel linear silicon probe in monocular V1 (Parker et al., 2022, 2023) . In both labs, we used KiloSort for spike sorting and Phy for curation (Lenzi & Steinmetz, 2020; Pachitariu, Pennington, et al., 2024; Pachitariu, Sridhar, et al., 2024).

The primary form of data with which ViSoND has been used is spiking data. The output files from spike sorting with Kilosort and Phy can be directly loaded into our pipeline that converts spiking to MIDI files. For other types of data, we must generate an event file, which has two columns: one column holds the event identities, which will each be matched to a particular pitch, while the second column holds the event times. This event file can be loaded into the pipeline.

### Conversion to MIDI

Our pipeline converts discrete event data into a MIDI format. Code is available in the Smear lab GitHub page (https://github.com/Smear-Lab/ViSoND). For our purposes, we treat a MIDI file as a six-column vector. The first column is the track, which designates the different event types for visualization and sonification. In one example, one track can be inhalation times mapped to a drum, while a second track can be spike times mapped to piano notes. The number of tracks used is up to the user. Other events can be included or different groups of neurons could be set to different tracks. The second column is the channel, which is a collection of tracks, which we set to a constant. The third column is the note, which can be any pitch from 0 to 127. Here, we restrict the pitches to a pentatonic scale to minimize dissonance, but that is not required. The fourth column holds a variable called velocity, which in music would essentially represent how hard you hit a piano key. We set velocity as a constant, but this could potentially be a useful parameter for ViSoND in future iterations. The fifth column holds the note onset time, in units of seconds on a grid with user-defined precision. We use a grid of 1e4 points per second. The sixth and final column is the note offset time, which we set to a constant (10 ms), although varying this could be useful in future iterations.

### Using the ViSoND

Once the MIDI file is complete, the next step is to play it with the video. The most basic way to do this on any operating system for free is to use VLC media player (https://www.videolan.org/vlc/index.html; VideoLan, 2006) . By installing an audio codec that can load MIDI files into VLC, as well as an accompanying SoundFont file (see GitHub for recommendations), the video and MIDI files can be played simultaneously.

For more flexible functions and data browsing, we worked with the ViSoND in Ableton Live (https://www.ableton.com/en/live/; Ableton, 2026) . Working in Live has the advantage of being able to zoom in and out of data with basic mouse gestures, and the ability to play back the data at a wide range of speeds. Many other digital audio workstations also enable alignment of MIDI and video, but we have yet to work with any of these. We have provided an Ableton Live template for ViSoND to allow for easy drag-and-drop upload and examination of files.

In videos 1 and 2, we mapped MIDI files based on spike times to notes played by a piano plug-in built into Live. In video 1, we rendered inhalation times to MIDI notes where pitch was mapped to log sniff rate. These MIDI files drove a kick drum plug-in built into Live. In video 2, we mapped gaze shift times to a custom handclap plug-in (https://www.maxforlive.com/library/device/1138/palmas; Clayton, 2012) . The user view of Ableton Live was screen captured with Open Broadcaster Software (https://obsproject.com/). Both ViSoNDS are played 3 times slower than real time.

**Video 1.**
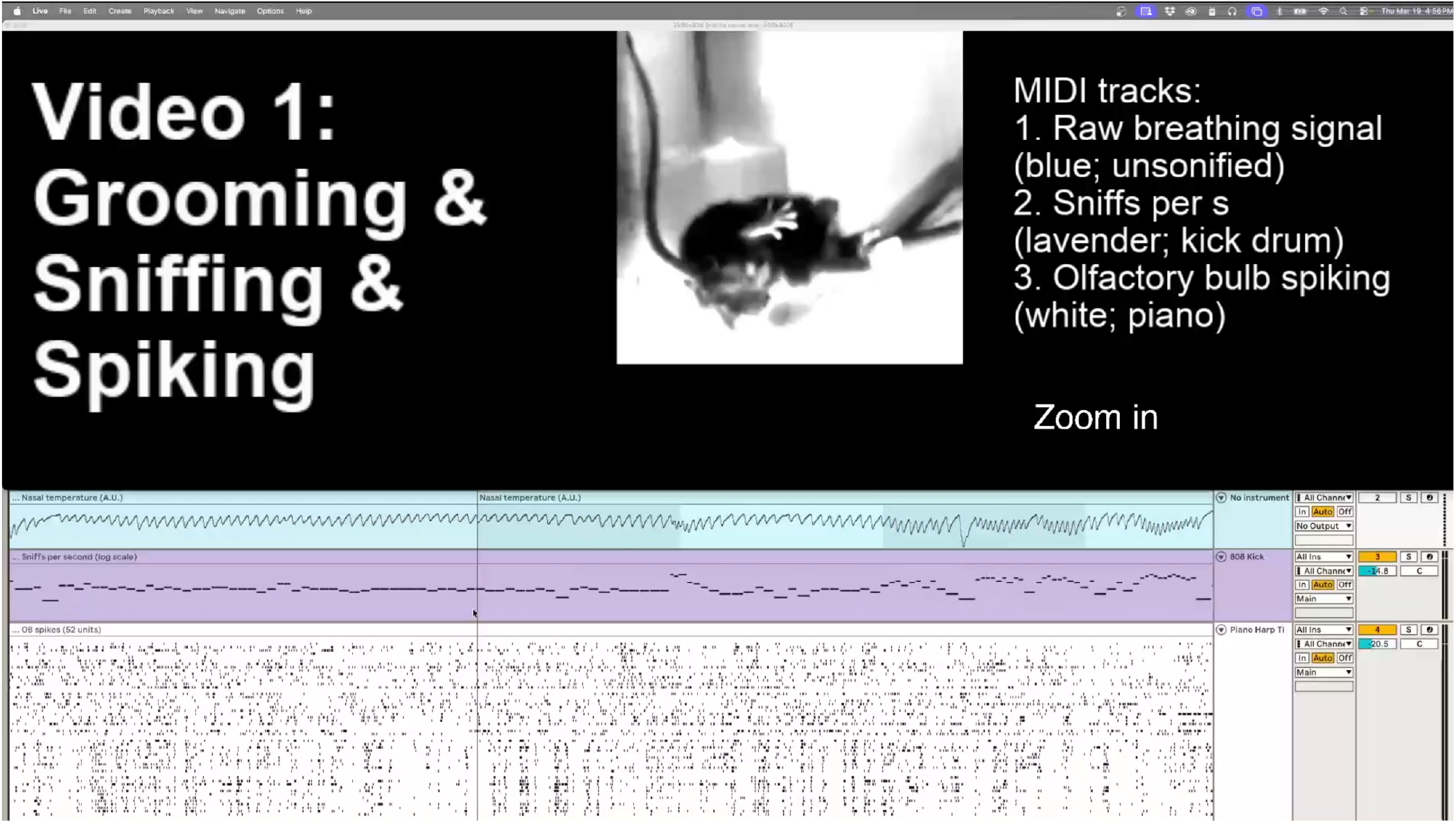
ViSoND reveals breathing-behavior association. This video shows that an intermediate-frequency breathing rhythm consistently accompanies grooming behavior. Using ViSoND, we zoom in from a full recording session (∼1 hour) to three ∼1-minute clips where this rhythm occurs. The display shows four panels: mouse video cropped and centered on the animal’s center of mass (top), raw breathing signal (blue), inhalation times with sniff rate (lavender), and olfactory bulb spikes from 52 neurons (white). Audio includes two sonified tracks: inhalation events (kick drum, pitch = sniff rate) and neuronal spikes (piano, one pitch per neuron). Video is played at 1/3 real-time speed. Zooming out and in transitions separate the three grooming examples. Observe that head, tongue, and paw movements often align with the breathing rhythm across all three bouts. This association was identified by navigating the sonified data, then validated by comparing multiple instances.

**Video 2.**
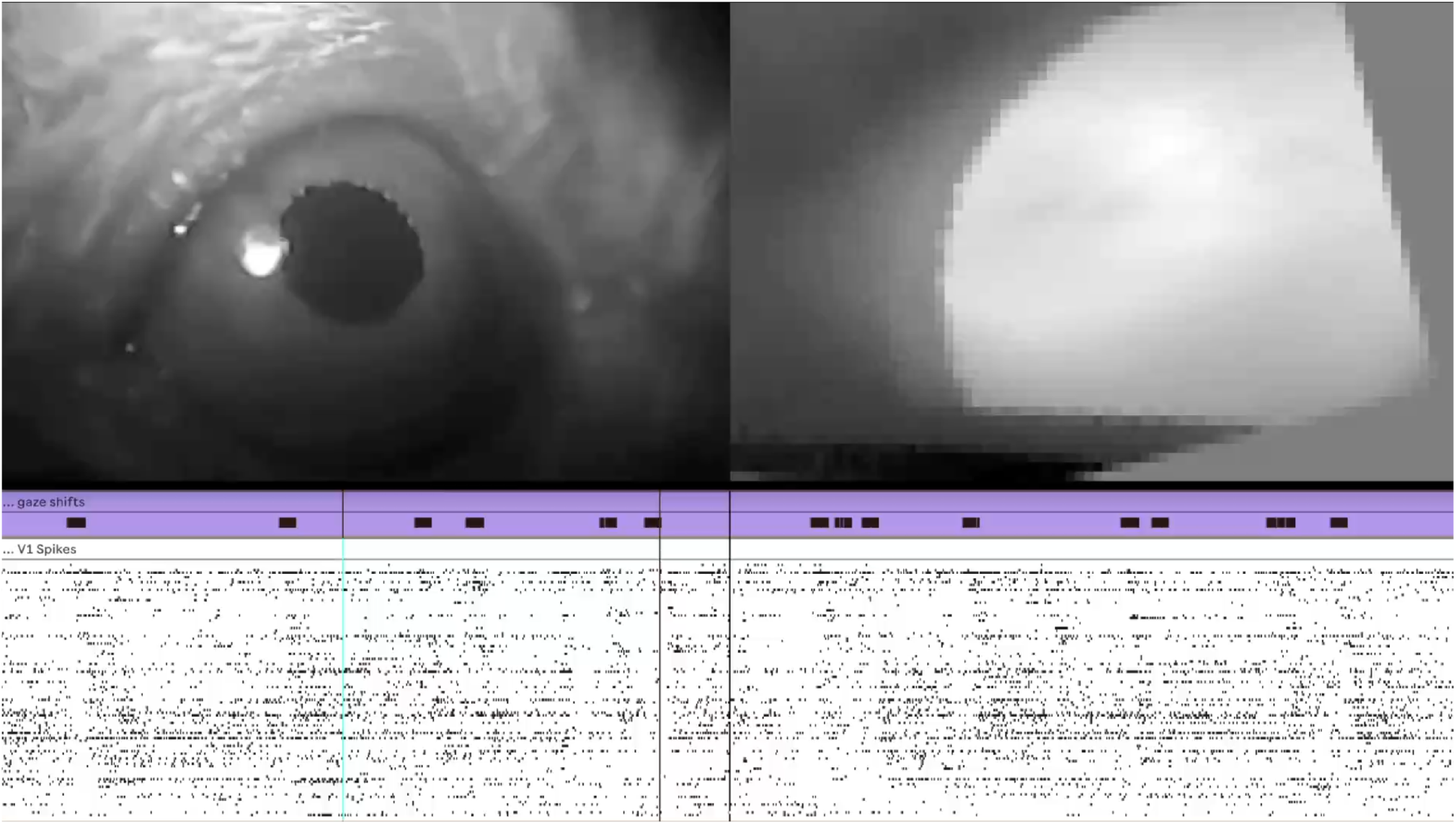
ViSoND reveals a blink response in mouse primary visual cortex. This video shows that blinks evoke sequential firing in V1, similar to the gaze-shift response. The top panel shows video from two cameras mounted on the mouse’s head: the left pane shows the eyecam view, and the right pane shows the worldcam view, adjusted according to a convolutional neural network (Parker et al., 2023). Below the video are two MIDI tracks in Ableton Live: gaze shifts (lavender; hand clap) and V1 spikes (white; piano, one pitch per neuron). Blinks are marked by vertical lines. Video is played at 1/3 real-time speed. Observe that spike sequences follow blinks with similar timing to gaze-shift responses. This blink response was not detected in the original analysis because pupil tracking failed during blinks. ViSoND enabled observation of these excluded periods, revealing that the blink response is qualitatively similar to the gaze-shift response.

## Results

Rigorous study of behavior requires precise quantitative measurements, but quantification necessarily entails reduction. The choices of what to quantify may not readily capture other relevant aspects of the biology. ViSoND provides a complementary approach to supplement quantification and computation with enhanced observation. Here we provide two use cases where ViSoND has yielded new insight into previously published datasets.

### A breathing rhythm identifies a grooming state in movement and spikes

Breathing synchronizes with behavior, and different breathing rates associate with different behavioral states (Sterrett et al., 2025). Furthermore, breathing synchronizes with neural activity in many areas of the brain (Heck et al., 2017; Karalis & Sirota, 2022; Tort et al., 2025; Zelano et al., 2016). To better understand the relationship between breathing and olfactory bulb activity, we have studied freely moving mice during ambient stimuli and uninstructed behavior. Even in this minimal condition, mice engage in highly structured breathing rhythms, which persist in a fairly stable frequency range for up to minutes at a time. These rhythmic states are paralleled by temporal structure in movement and olfactory bulb activity (Sterrett et al., 2025). Computational modeling identified an intermediate frequency breathing state that we had not anticipated and did not know how to interpret. Could this rhythmic state have an association with any particular form of behavior? Although computational methods identified interpretable behavioral structure in kinematic variables, we found that ViSoND was a more direct path to insight.

The key was rendering the breathing rhythm into MIDI and then visually navigating to periods in the video during which the mouse was using the intermediate-frequency breathing state (Video 1).

Observation indicates that this rhythmic state consistently accompanies grooming behaviors, where the visible movements of the tongue and paws synchronize in lockstep with the audible breathing rhythm. Further, this behavioral state co-occurs with a characteristic pattern of “spontaneous” activity of olfactory bulb neurons. With ViSoND, it is possible to quickly navigate between multiple instances of the rhythm of interest, and check whether such an audio-visual association is a one-off coincidence or if the intermediate breathing rhythm and grooming co-occur reliably. By concatenating video clips of the putative “grooming rhythm”, we show that this co-occurrence is frequent (Video 1 shows three examples). So, while this association between breathing and grooming could have been extracted with computational tools, ViSoND makes the relationship immediately obvious to eye and ear. This example illustrates how ViSoND can unite quantitative modeling with direct observation.

### A blink response in primary visual cortex

In an unrestrained animal, the head and eyes are almost always in motion. To study cortical processing during natural locomotion, we performed visual physiology in unrestrained mice allowed to move freely through a visually complex environment without a task or instruction (Fig 3A; Parker et al., 2022, 2023). During free movement, mice made frequent gaze-shifting saccadic eye movements which let the visual field slip across the retina, unlike compensatory eye/head movements which stabilize the retinal image (Fig 3B). The onset of each gaze shift evoked a temporal sequence of action potentials in mouse primary visual cortex (Fig 3B; (Parker et al., 2023). Neurons that prefer low spatial frequency tend to fire earlier, while those that prefer high spatial frequency tend to fire later, a pattern consistent with coarse-to-fine processing.

**Figure 3.**
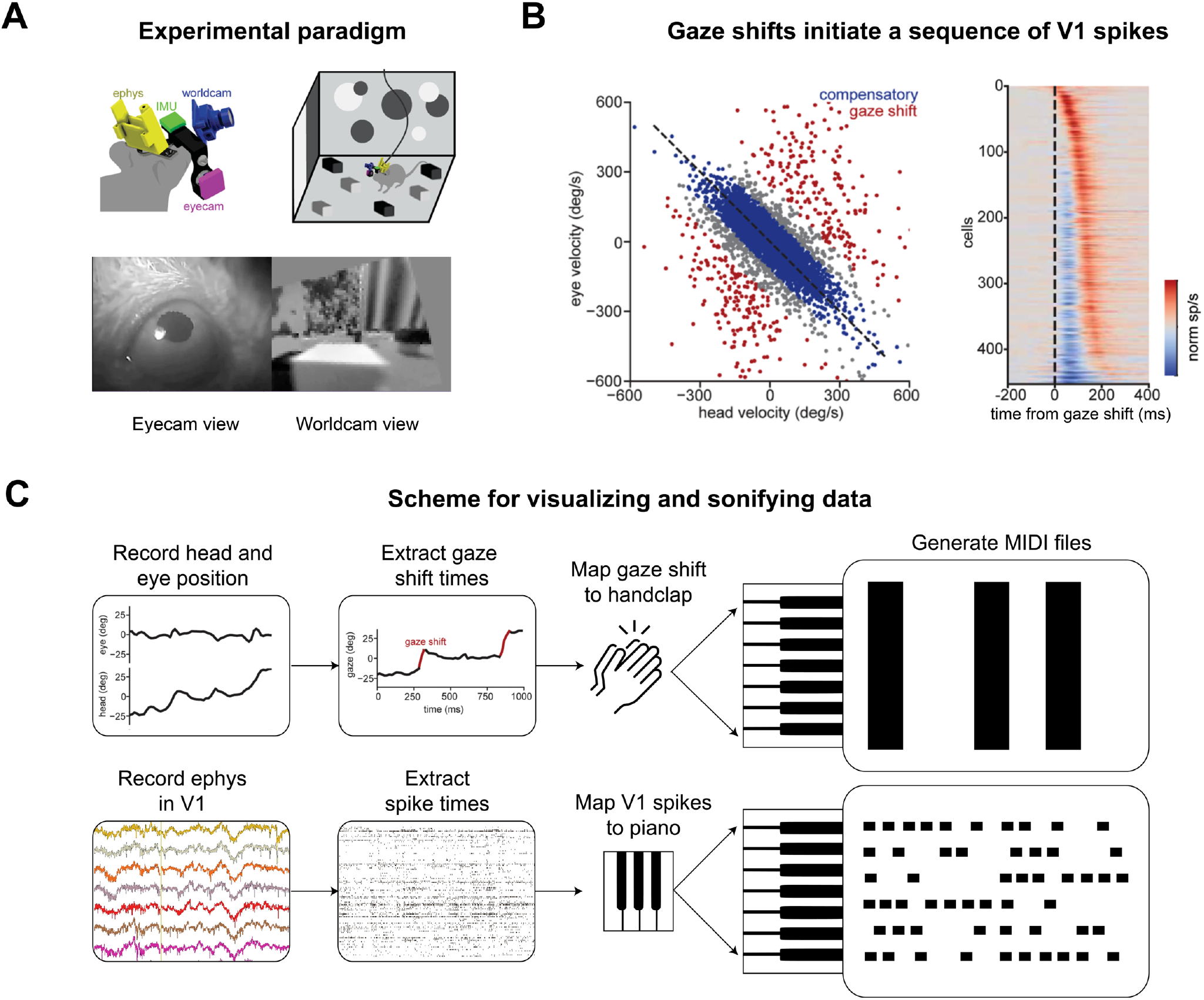
A blink response in primary visual cortex (accompanies Video 2). (A) Diagram of experiment with simultaneous recording of head movement, eye position, visual scene, and primary visual cortex (V1) activity in freely behaving mice. (B) *Left*. Gaze shifts are defined as events where a fast head movement is not counteracted by compensatory eye movement (data from one session). *Right*. Gaze shifts evoke sequential firing events in V1 population activity (n=9 mice) (C) Diagram of ViSoND pipeline for the experiment. Data in A and B adapted from Parker et al., 2023.

Our analysis of gaze shifts used pupil tracking to calculate eye direction, so we excluded times when the video frames lacked a usable image of the pupil. With ViSoND, we need not impose this curation, allowing us to observe that neural patterns of sequential firing accompanying gaze shifts also occur following a blink (Video 2). In this example as above, the relationship between blinking and V1 spikes could have been discovered with a different choice of quantification, but ViSoND made it possible to see and hear the relationship in action.

## Discussion

ViSoND is intended to enhance an observer’s ability to perceive patterns in data that may not be immediately apparent from quantitative, computational analyses. Sonification is deeply rooted in neurophysiological tradition going back to Lord Adrian (E. Adrian, 1932; E. D. Adrian & Bronk, 1928) . Using MIDI allows us to extend this tradition in a way that allows behavioral and neural dynamics to be seen, heard, and aligned with video from behaving animals. Additionally, the use of musical sound may lower the barrier to engaging the public with complex neural data in educational and outreach contexts.

Once we convert data to MIDI format, we can work with them in digital audio workstations (DAWs; e.g., Ableton Live), which have been optimized by professional coders to serve a thriving market of amateur and professional musicians. Our pipeline is simple to implement, requiring no previous experience working with MIDI or DAWs. Being able to observe neurophysiological data through multiple sensory modalities opens up new opportunities for data analysis and data communication.

For future directions, we see the potential to adapt ViSoND for other kinds of data. If fluorescence or BOLD traces can be discretized in a useful way, ViSoND may also facilitate pattern detection in calcium imaging or fMRI. Other kinds of data could also be dimensionally reduced for use with ViSoND, such as behavior structure, for example where certain behavior syllables are pitched to notes. Similarly, when the number of recorded neurons exceeds human auditory cognition by requiring “too many notes” (Hampe, 2016), ViSoND could render dimensionality-reduced spiking data in a useful way. More broadly, any data that can be expressed as a time series of discrete events can readily be converted to a ViSoND using code available on our Github page. We anticipate that future ViSoND users who are familiar with music theory and digital music software could innovate new ways of utilizing this technique and improve how well data can be represented through sonification. We hope that ViSoND can serve as a bridge between computational analysis of high-dimensional data and the qualitative, observation-driven insights that have historically driven neuroscience forward.

## Supporting information

Video 1 - A breathing rhythm associated with grooming

Video 2 - A blink response in primary visual cortex

## Acknowledgements

We thank Sid Rafilson, Emmalyn Leonard, Adrienne Fairhall, and James Murray for contributions to the published work, and members of the Smear & Niell labs for helpful conversations and feedback on the manuscript. This work was supported by NIH grants R01NS123903 (MCS), R01DC018789 (MCS), and the Simons Collaboration on the Global Brain (SCS), UF1NS116377 (CMN), and R01NS121919-01 (CMN). This research was supported in part by grant NSF PHY-2309135 to the Kavli Institute for Theoretical Physics (KITP).

